# ZapC crosslinks FtsZ filaments through a dual-binding mechanism modulated by the intrinsically disordered linker of FtsZ

**DOI:** 10.1101/2025.05.07.652675

**Authors:** Ying Li, Han Gong, Rui Zhan, Yuanyuan Cui, Xiangdong Chen, Joe Lutkenhaus, Shishen Du

## Abstract

Most bacteria divide through binary fission, which is mediated by a large protein complex called the divisome. Assembly of the divisome is initiated by formation a Z ring at midcell consisting of polymers of the bacterial tubulin FtsZ. A series of FtsZ-associated proteins (Zaps), which crosslink FtsZ filaments, promote Z ring formation in *Escherichia coli*. However, how these proteins interact with FtsZ is still unclear. In this study, we discover that ZapC binds to both FtsZ’s globular domain and its conserved C-terminal peptide (CCTP) to crosslink FtsZ filaments. An AlphaFold 3 structural model of the FtsZ-ZapC complex indicates that ZapC binds to the globular domain of FtsZ via a loop region connecting its N-terminal and C-terminal domains and to the CCTP of FtsZ via a hydrophobic pocket in the N-terminal domain. Substitutions in these regions of ZapC disrupt its binding to FtsZ, validating the dual binding mode. Strikingly, we find that the intrinsically disordered C-terminal linker (CTL) of FtsZ affects the interaction of FtsZ with ZapC as well as other partners, indicating an important role of the CTL in FtsZ functionality. Taken together, these results indicate that ZapC, although it exists as a monomer, can crosslink FtsZ filaments by a two-pronged mechanism, binding to the globular domain of FtsZ in one filament and to the CCTP of FtsZ in another filament. Furthermore, the CTL plays an important role in regulating FtsZ interaction with its partners.

**Importance:** Bacterial cytokinesis requires the Z ring, a highly dynamic cytoskeletal element consisting of polymers of the bacterial tubulin FtsZ. Formation of a coherent and functional Z ring is facilitated by FtsZ-associated proteins (Zap), which can crosslink FtsZ polymers, but how these proteins work is still incompletely understood. In this study, we find that ZapC, one of the FtsZ crosslinkers, binds to both FtsZ’s globular domain and its conserved C-terminal peptide to crosslink FtsZ filaments. Moreover, the intrinsically disordered C-terminal linker (CTL) of FtsZ modulates its binding to ZapC and many other FtsZ binding proteins. These findings reveal a novel mechanism to crosslink FtsZ filaments and reveal an important and highly conserved role of the CTL in FtsZ functionality.

## Introduction

During bacterial cytokinesis, the tubulin-like protein FtsZ assembles into the Z ring at midcell to recruit the other division proteins to assemble the division apparatus (divisome), which synthesizes the septum, leading to the generation of two daughter cells (1–3). FtsZ is composed of four structural domains: a short N-terminal motif, a globular domain with GTPase activity, an intrinsically disordered C-terminal linker (CTL) of variable length, followed by a highly conserved C-terminal peptide (CCTP) (4, 5). The globular domain of FtsZ binds to GTP and polymerizes into protofilaments which depolymerize following GTP hydrolysis (6). At steady state, FtsZ protofilaments undergo treadmilling, in which subunits are added to one end of filaments and leave from the opposite end (5, 7–9). The CCTP of FtsZ mediates its interactions with membrane anchors (FtsA and ZipA in *E. coli*), thus tethering FtsZ filaments to the membrane (10, 11). The CCTP also mediates binding to proteins that regulate FtsZ function, including both positive and negative regulators. For example, MinC and SlmA, which are components of the Min and Nucleoid Occlusion systems regulating Z ring positioning, respectively, grasp the CCTP to bind to FtsZ filaments and antagonize its polymerization (12–14). Although the CTL is not directly involved in FtsZ polymerization, recent studies found that its length and flexibility are important for proper FtsZ function (15–21). Moreover, the CTL may also affect the interaction between FtsZ and its modulatory proteins (18). The extreme N-terminal motif of FtsZ is less studied, but recent studies found that it is widely conserved and involved in interaction with ZapA, an FtsZ-associated protein (22, 23).

Assembly of a coherent functional Z ring is a premise for successful cell division because the Z ring not only defines the site for division, but also serves as the scaffold for assembly of the divisome complex (2, 4, 5). The Z ring is also believed to provide the force for membrane deformation during constriction initiation (4, 5). Also, the treadmilling dynamics of the FtsZ filaments in the Z ring help to distribute the septal peptidoglycan synthetic complex FtsQLBWI along the circumference to build a smooth septum (7, 8, 24–26). Due to the crucial role of the Z ring in bacterial cytokinesis, its assembly, dynamics and stability have been extensively investigated. Recent studies showed that FtsZ filaments initially form a loose structure on the membrane at the midcell, which subsequently condenses into a coherent Z-ring with the help of FtsZ-associated proteins (Zaps) also called FtsZ-binding proteins (ZBPs)(26, 27). In *E. coli*, the identified Zap proteins thus far include ZapA, ZapB, ZapC, ZapD and ZapE (28–34). Most Zap proteins, with the exception of ZapB, interact directly with FtsZ to cross-link FtsZ filaments, thus promoting the condensation of FtsZ protofilaments into the Z ring (34). However, all known Zap proteins are nonessential for division and their absence typically results in minor or modest abnormalities in Z-ring structure and– septal morphology (27, 34). Nonetheless, when multiple Zap proteins are absent, FtsZ filaments fail to condense into a stable and tightly-ordered Z-ring, leading to significant defects in cell division, indicating that they work together to facilitate Z ring assembly and stability (27, 34).

The molecular mechanisms through which Zap proteins promote Z-ring formation have been extensively studied. ZapA is a widely conserved protein present in almost all bacteria (28). ZapA primarily exists as tetramers within the cell which crosslink and align FtsZ filaments into the Z ring (35–39). Recent studies showed that a ZapA tetramer grabs the N-terminal tail of FtsZ and binds to the junctions between FtsZ subunits within filaments to crosslink and align them in a parallel fashion (23). Unlike ZapA, ZapB does not bind directly to FtsZ. Instead, it is recruited to the division site through an interaction with ZapA (40) and MatP, which is an organizer of the Ter macrodomain of the chromosome (41, 42). Thus, ZapA, ZapB and MatP form a multilayer protein network termed the Ter linkage that facilitates rapid Z ring assembly and coordinates cell division with chromosome segregation (42–44). The working mechanism of ZapD is most clearly understood, it forms dimers that interact with the CCTP of FtsZ to crosslink FtsZ filaments (45).

ZapC can also crosslink FtsZ filaments into large bundles *in vitro* (30, 31), but how it works has been controversial. Bhattacharya et al. found that ZapC’s binding to FtsZ differs significantly from that of ZipA which binds the CCTP and thus concluded that ZapC does not interact with the CCTP (46). In agreement with this, Schumacher’s team found through yeast two-hybrid and sedimentation assays that the CCTP of FtsZ was not critical for its interaction with ZapC (47). However, Ortiz et al. detected an interaction between the CCTP of FtsZ and ZapC using Isothermal Titration Calorimetry (ITC), albeit with extremely low affinity (48). The crystal structure of ZapC revealed that it exists as a monomer composed of two domains, each of which possesses a binding pocket (47). Mutations in or near these two pockets disrupt the function of ZapC and prevent it from binding to FtsZ. Thus, it was proposed that these pockets bind to the globular domain of FtsZ subunits in adjacent FtsZ filaments, thereby facilitating their cross-linking (47). However, whether this model is correct remains unknown. Given that ZapC is a monomer, the mechanism by which it crosslinks FtsZ filaments is likely distinct from the other Zap proteins, such as ZapA and ZapD, which exist as multimers (tetramer or dimer) and bind the N-terminal tail or CCTP (35, 39, 45). Thus, further investigation of the interaction between ZapC and FtsZ may not only resolve the controversy regarding its working mechanism, but also uncover novel mechanisms for regulating FtsZ assembly.

In this study, we characterized mutations affecting the interaction between FtsZ and ZapC through genetic, biochemical and cellular approaches. Our results indicate that ZapC monomers bind to both the globular domain and the CCTP of FtsZ to crosslink FtsZ filaments. Moreover, the CTL of FtsZ can significantly affect the binding of the CCTP to ZapC, as well as other CCTP binding proteins, providing direct evidence for the modulation of FtsZ’s interaction with its binding partners by the CTL.

## Results

### Identification of FtsZ mutations providing resistance to ZapC overexpression toxicity

Overexpression of ZapC blocks Z-ring formation resulting in cell division inhibition and cell death (30, 31). Although mutations in ZapC that prevent cell death have been isolated and used to identify two pockets in ZapC likely to be involved in binding FtsZ, mutations in FtsZ that prevent cell death have not been identified. Isolation and study of such mutations may reveal details of the interaction mechanism between them. Such an approach was very successful in elucidating details of the interaction of MinC/MinD and SlmA with FtsZ (12, 13, 49). Therefore, we constructed an FtsZ mutant library and screened for *ftsZ* mutations that conferred resistance to ZapC overexpression toxicity as illustrated in Fig. S1A and described in the Materials and Methods. Cells expressing wild-type FtsZ could not form colonies on plates containing 15 or more μM IPTG to induce the expression of ZapC (Fig. S1). However, 13 mutant strains grew on plates with 60 μM IPTG, suggesting that they suppress the toxicity of ZapC. Sequencing of the *ftsZ* coding regions from these 13 mutants revealed that they contained single or multiple mutations in *ftsZ* (Table S1). Subsequent analysis revealed that a total of 10 mutations in FtsZ exhibit varying degrees of resistance to ZapC overexpression (Fig. 1A and Table S2). Among them, F285S、E322K and I323L displayed the strongest resistance, followed by F377Y, A376T and K380M, while E147G, N359Y and D360Y provided only weak resistance (Fig. 1A). Examination of cell morphologies showed that cells expressing wild-type FtsZ were extremely filamentous following induction of ZapC with 30 μM IPTG, however, cells expressing the FtsZ mutants were much shorter under the same condition (Fig. 1B), implying that they provide resistance to the action of ZapC.

**Figure 1.**
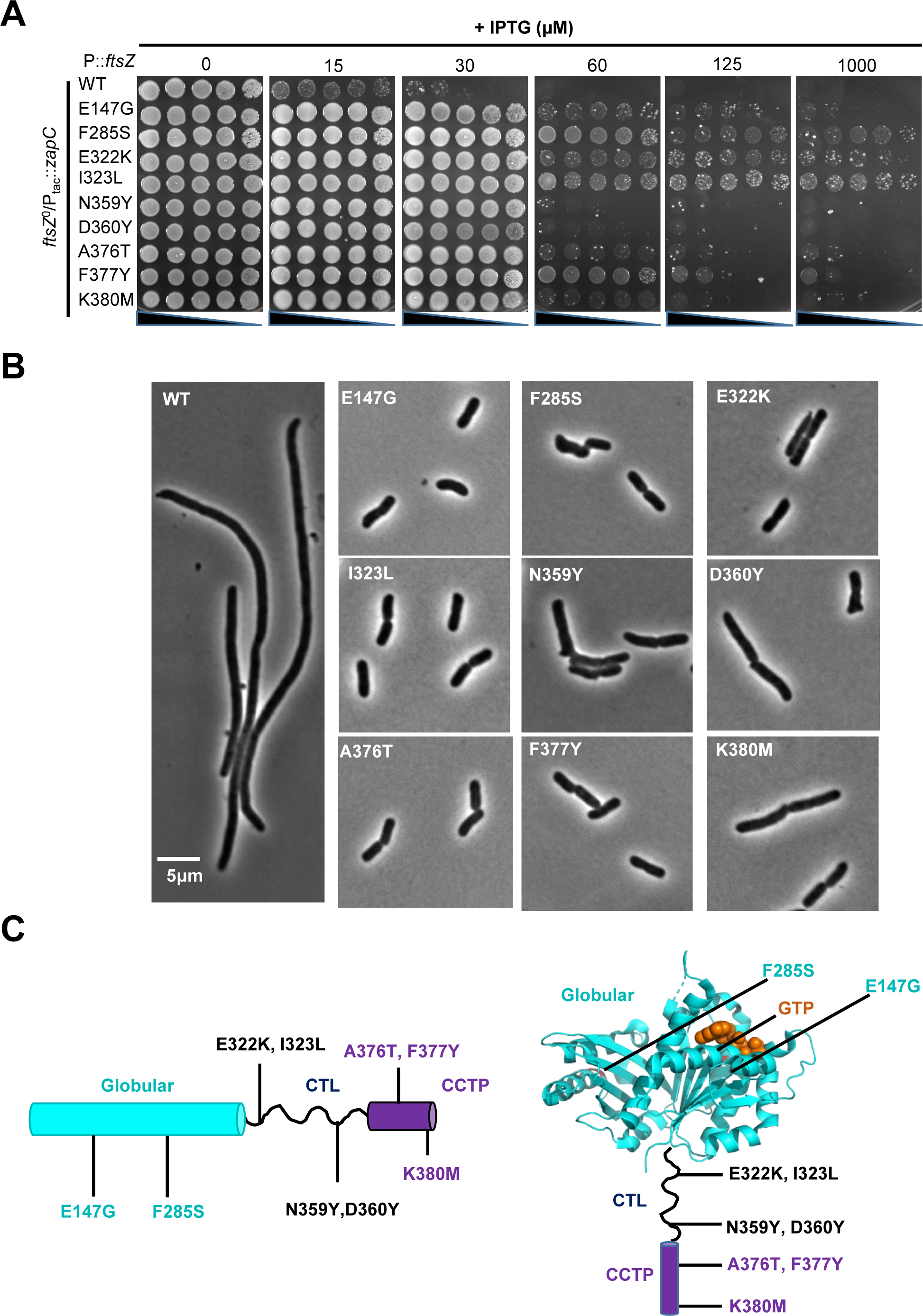
Identification of FtsZ mutations providing resistance to ZapC overexpression toxicity. (A) A spot test of FtsZ mutants displaying resistance to ZapC overexpression toxicity. Plasmids pBANG112 (p15A, *ftsZ*) or its derivatives carrying different *ftsZ* alleles were transformed into strain S17/pKD3C (W3110, *ftsZ*0 /pSC101^ts^, *ftsZ*) harboring plasmid pSD320 (pEXT22, P_tac_::*zapC*) and transformants were incubated at 42°C with antibiotics. The next day, a single colony of each resulting strain was subject to spot tests on LB plates with different concentrations of IPTG. Plates were incubated at 37°C overnight and photographed. (B) Representative images of morphology of cells expressing FtsZ or its variants and overexpressing ZapC from 3 biological replicates. Strains were grown at 37 °C in exponential phase, FtsZ mutants were constitutively expressed from plasmid pBANG112 and ZapC was induced with 30 μM IPTG for 3 h before imaging. Scale bar, 5 μm. (C) Location of ZapC-resistant mutations in FtsZ. Mutations in the globular domian (E147G and F285S), intrinsically disordered linker (E322K, I323L, N359Y and D360Y) and conserved C-terminal peptide (A376T, F377Y and K380M), are indicated, and colored cyan, black and purple in the topology and structure of FtsZ (PDB#: 6UNX), respectively.

### ZapC resistant mutations are located in the globular domain, CTL and CCTP of FtsZ

Mapping the ZapC resistant FtsZ mutations onto the primary sequence of *E. coli* FtsZ (PDB#: 6UNX)(50) revealed that they are scattered along the length of the protein, with E147G and F285S residing in the globular domain, E322K, I323L, N359Y, and D360Y in the CTL, and A376T, F377Y and K380M in the CCTP (Fig. 1C). This suggested that multiple domains of FtsZ are involved in the binding to ZapC and a combination of these mutations might provide greater resistance to ZapC overexpression. To test this, we selected a mutation from each domain and constructed double and triple mutants of FtsZ. Complementation tests of these double or triple mutants showed that they could complement an FtsZ depletion strain, suggesting that they retained the basic function of FtsZ (Fig. S2A). As expected, these double or triple mutations of FtsZ, except for I323L and D360Y, displayed stronger resistance to the toxicity of ZapC overexpression in comparison to their respective single mutations, indicating that they have additive effects (Fig. S2B). Notably, while each of the single mutations in the CTL or CCTP of FtsZ provided only weak or modest resistance to ZapC overexpression, the triple mutant (I323L, D360Y and A376T) allowed the cells to grow on plates with even 1 mM IPTG. This implies that the globular domain, the CTL and the CCTP of FtsZ are all directly or indirectly involved in the interaction with ZapC.

### ZapC resistant FtsZ mutations prevent the midcell localization of ZapC

Previous studies have shown that mutations in ZapC, which disrupt its interaction with FtsZ, prevent it from localizing to the midcell (47). If the FtsZ mutations disrupt the interaction between FtsZ and ZapC, they should also block the localization of ZapC. Thus, we fused ZapC to GFP and examined its localization in cells expressing the FtsZ mutants (note that ZapC-GFP was expressed at a level that did not block Z ring formation in these experiments). To ensure that the FtsZ mutations did not substantially affect the assembly of the Z-ring, we also examined the localization of ZapA-mCherry, which serves as a marker for the Z ring. As shown in Fig. 2A and Table S3, ZapA-mCherry localized as a band in cells expressing the FtsZ mutants, suggesting that they retained the ability to form functional Z rings and the mutations did not affect the interaction with ZapA. As expected, ZapC-GFP co-localized with ZapA-mCherry as a sharp band in 98.8% of the cells expressing wild-type FtsZ. However, ZapC-GFP exhibited different degrees of localization defects in cells expressing the single FtsZ mutants. In most cells expressing FtsZ^F285S^, ZapC-GFP was evenly distributed in the cytoplasm. As a consequence, its co-localization with ZapA-mCherry dropped to only 8.5%, indicating that F285S greatly reduced the interaction between FtsZ and ZapC. In cells expressing FtsZ^A376T^, the co-localization of ZapC-GFP with ZapA-mCherry dropped to about 44%, indicating that it modestly reduced the FtsZ interaction with ZapC. Interestingly, although the mutations in the CTL of FtsZ (I323L or D360Y) provided resistance to ZapC overexpression as shown above, ZapC-GFP co-localized very well with ZapA-mCherry in cells expressing the respective FtsZ mutants, suggesting that these mutations did not significantly disrupt the interaction. The co-localization of ZapC-GFP and ZapA-mCherry was greatly reduced in cells expressing the double or triple FtsZ mutants, except for the double mutant containing the I323L and D360Y mutations, both of which are located in the CTL. However, when either one of them was combined with F285S, the midcell localization of ZapC-GFP was further reduced in comparison to that of F285S alone. Also, the combination of I323L and A376T, which are in the CTL and the CCTP, respectively, reduced the co-localization of ZapC-GFP and ZapA-mCherry to about 16%. Taken together, these results indicate that the mutations in the CTL also negatively affect the localization of ZapC-GFP, but their effect only becomes evident when they are combined with mutations in the globular domain or the CCTP.

**Figure 2.**
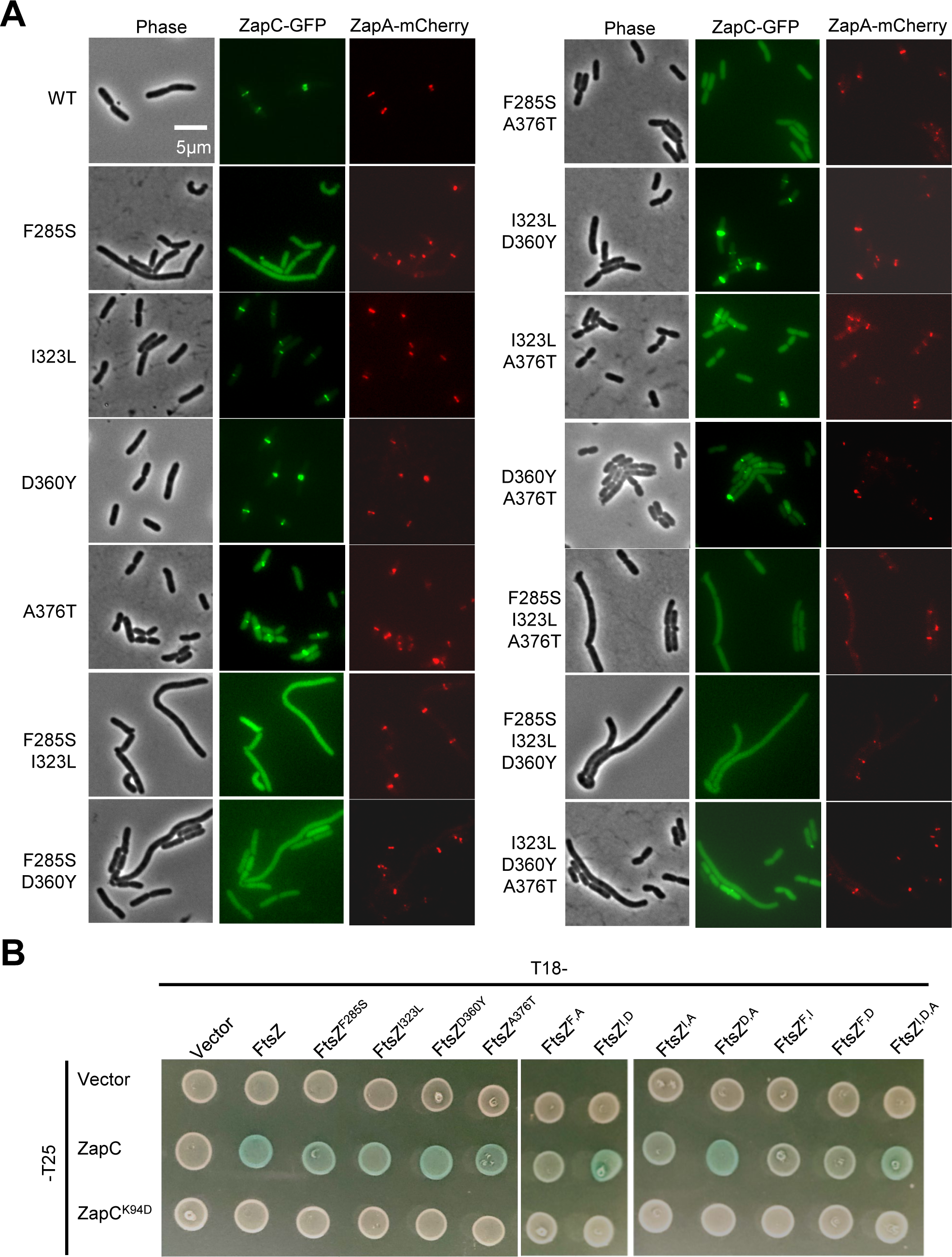
ZapC-resistant mutations weaken the interaction between FtsZ and ZapC *in vivo*. (A) Representative images of ZapC-GFP and ZapA-mCherry localization in cells expressing FtsZ or its mutants. Overnight cultures of cells expressing wild type FtsZ or its variants and ZapC-GFP and ZapA-mCherry were diluted 1:100 in fresh LB medium with antibiotics and cultured at 37°C. 3 hours later the cultures were diluted 1:10, IPTG was added to a final centration of 50 μM to induce the expression of ZapC-GFP. 2.5 hours post induction cells were immobilized on 2% agarose pads for photographing. ZapA-mCherry was constitutively expressed from its chromosomal locus. Scale bar, 5 μm. (B) Bacterial two hybrid assay to test the interaction between FtsZ or its mutants and ZapC or ZapC^K94D^. Plasmids pairs were transformed into strain LYA1 (BTH101, *ftsA*^R286W^), the next day a single transformant of each resulting strain was resupended in 1 mL LB medium and 2 μL of each culture was spotted on LB plates containing antibiotics, 40 μg/mL X-gal and IPTG. Plates were incubated at 30°C for about 18 hours before photographing.

### ZapC resistant FtsZ mutations reduce the interaction between FtsZ and ZapC *in vivo*

To confirm that the FtsZ mutations weaken the interaction between FtsZ and ZapC *in vivo*, we checked their impact on the interaction using the bacterial two hybrid (BTH) assay (51). Initially, we used the specialized strain BTH101 for the experiment. However, we found that the strain was not suitable for testing the interaction between FtsZ and ZapC because transformants carrying the plasmids expressing the fusions grew very poorly even without induction, likely due to the toxicity of these fusions. To overcome this problem, we introduced the *ftsA\** (FtsA^R286W^) mutation into the BTH101 strain, since FtsA* interacts more strongly with FtsZ and has been reported to reduce the toxicity of ZapC overexpression (48, 52, 53). We confirmed that the resultant strain (LYA1) containing *ftsA\** provided modest resistance to overexpression of ZapC and allowed the transformation of the plasmid pairs into the strain (Fig. S3A). As expected, a clear interaction signal between ZapC and FtsZ (colonies turning blue) was observed, as well as with FtsZ itself (Fig. 2B and Fig. S3B). Introduction of the K94D mutation into ZapC, which has been reported to disrupt ZapC interaction with FtsZ (47), eliminated the interaction signal (Fig. 2B), suggesting that the assay could be used to access the interaction between FtsZ and ZapC.

Unexpectedly, introduction of the single mutations into the FtsZ-T25 fusion protein did not significantly reduce its interaction with ZapC (Fig. 2B). The interaction signal between FtsZ^F285S^ or FtsZ^I323L^ and ZapC was only slightly reduced compared to that between wild-type FtsZ and ZapC (Fig. 2B), while D360Y and A376T did not detectably reduce the interaction signal. However, combinations of these mutations significantly reduced or completely eliminated the interaction signal. For example, a combination of F285S with any of the other three mutations eliminated its interaction with ZapC. Also, the interaction signal between FtsZ and ZapC disappeared when A376T was combined with I323L. These results indicate that all ZapC resistant FtsZ mutations affected the interaction, but not all are disruptive enough to produce an obvious reduction in the interaction signal in the BTH assay, unless they were combined. These results are consistent with those of the toxicity test and the localization study, which showed that the effect of the mutations were additive. Lastly, we found that none of the FtsZ mutations affected the interaction between FtsZ and ZapA (Fig. S3C), suggesting that the impact of the mutations was specific to ZapC. Taken together, these results confirm that the ZapC resistant FtsZ mutations weaken the interaction between FtsZ and ZapC *in vivo*.

### ZapC resistant FtsZ mutations reduce the interaction between ZapC and FtsZ *in vitro*

To confirm that the ZapC resistant FtsZ mutations disrupt the interaction between FtsZ and ZapC, we purified ZapC, FtsZ and their variants using the SUMO-tag purification system and tested their interaction *in vitro*. We first ensured that the FtsZ mutants could polymerize *in vitro* using a sedimentation assay. As shown in Fig. S4, all FtsZ mutant proteins, similar to wild-type FtsZ, sedimented to the pellet after ultrahigh speed centrifugation in the presence of GTP and Ca^2+^ but not in the presence of GDP, indicating that they could polymerize into filaments. Note that the amount of FtsZ^F285S^ in the pellet was relatively less compared to wild-type FtsZ, suggesting that the mutation somehow affects assembly. The addition of ZapC to the reaction greatly increased the amount of wild-type FtsZ in the pellet (in the absence of Ca^2+^), and ZapC was also deposited to the pellet (Fig. 3A), suggesting ZapC promotes the formation of FtsZ filament bundles. Interestingly, ZapC could still promote the sedimentation of the single FtsZ mutants, despite that they interacted with ZapC less well *in vivo*. Nonetheless, ZapC could not promote the sedimentation of the double mutant FtsZ^F285S, A376T^, which contains a mutation in the globular domain and another in the CCTP (Fig. 3A). This indicates that the combination of these two mutations significantly weakens the interaction between FtsZ and ZapC *in vitro*.

**Figure 3.**
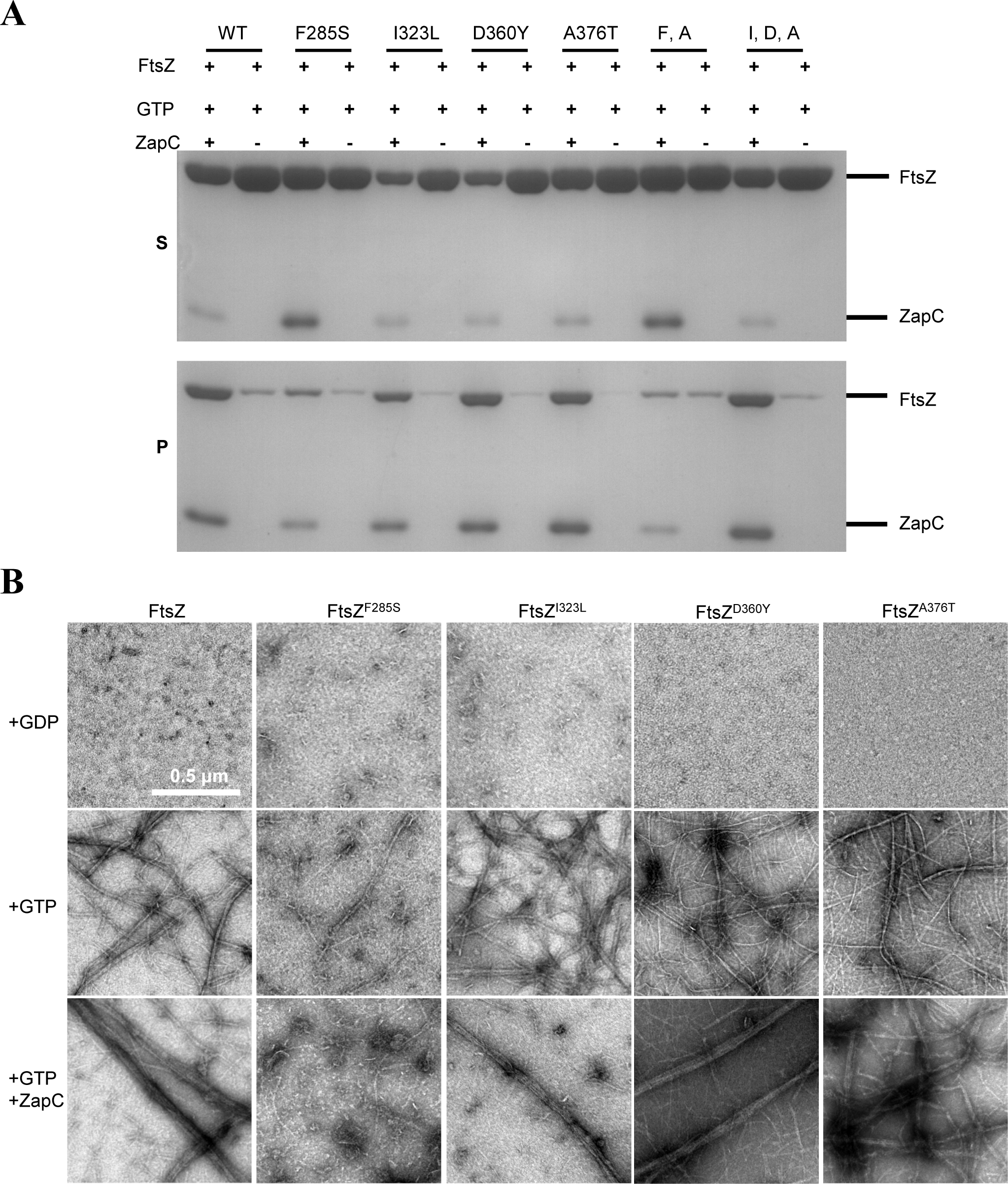
ZapC-resistant mutations weaken the interaction between FtsZ and ZapC *in vitro*. (A) Co-sedimentation assay to assess the impact of ZapC on FtsZ polymerization. FtsZ and 6×His-ZapC were added at a final concentration of 5 μM in a total polymerization reaction volume of 50 μL. The reactions were carried out as described in Materials and Methods. The amount of FtsZ in the supernatant (S) and pellet (P) was analyzed by SDS-PAGE. (B) FtsZ mutants are resistant to the crosslinking activity of ZapC. Polymerization of FtsZ was performed as in the co-sedimentation assay and samples for transmission electron microscopy were prepared as described in Materials and Methods. Representative images of FtsZ in the presence of GDP, GTP, or GTP and 6×His-ZapC are shown. Double and triple mutants are indicated by the capital letters of the mutated residues. Scale bar, 0.5 μm.

Examination of the polymers formed by FtsZ or its mutants using negative stain electron microscopy showed that all of them assembled into single or double filaments in the presence of GTP but not GDP, confirming they polymerized into filaments. However, FtsZ^F285S^ displayed less filaments on the grid, consistent with its reduced ability to assemble as indicated by the sedimentation assay. The addition of ZapC to wild-type FtsZ resulted in the formation of large filament bundles as previously reported (Fig. 3B). However, its effect on the FtsZ^F285S^ filaments was negligible, indicating that ZapC could not effectively crosslink filaments formed by FtsZ^F285S^. Interestingly, although ZapC could facilitate the sedimentation of FtsZ mutants harboring the I323L, D360Y, or A376T mutations, we observed much fewer filament bundles in comparison to wild-type FtsZ, especially for the mutants containing the A376T mutation (Fig. 3B). Thus, the FtsZ mutants show varying degrees of resistance to the cross-linking activity of ZapC *in vitro*.

To quantify the impact of these mutations on the interaction between FtsZ and ZapC, we measured the binding affinity between them using biolayer interferometry. We attached His-tagged ZapC protein to the biosensor and then monitored the binding of FtsZ or its mutants. As shown in Fig. 4A-E, wild-type FtsZ bound to ZapC with fast kinetics and dissociated rapidly with a dissociation constant (Kd) of 0.39 μM, indicating that FtsZ binds ZapC with a high affinity. All the single FtsZ mutants (F285S, I323L, D360Y and A376T) bound to ZapC, but the Kd values were reduced significantly compared to wild-type FtsZ. FtsZ mutants harboring mutations in the CTL or CCTP (I323L, D360Y and A376T) displayed Kd values 1.5-2 times higher than that of wild-type FtsZ (Table S4), whereas FtsZ^F285S^ showed a 16-fold reduced Kd (0.39 μM vs 6.37 μM), indicating that the former group of mutations modestly weaken the interaction, while F285S has a more drastic effect on the interaction. Moreover, FtsZ^F285S^ displayed different binding kinetics to ZapC as ZapC bound slowly and did not dissociate in the dissociation step (Fig. 4B). This suggests this mutant binds to ZapC by a different binding mode in comparison to wild-type FtsZ. Taken together, these results indicate that the FtsZ mutations reduce FtsZ’s ability to interact with ZapC to different extents, consistent with the varied resistance to ZapC overexpression and ZapC localization *in vivo*.

**Figure 4.**
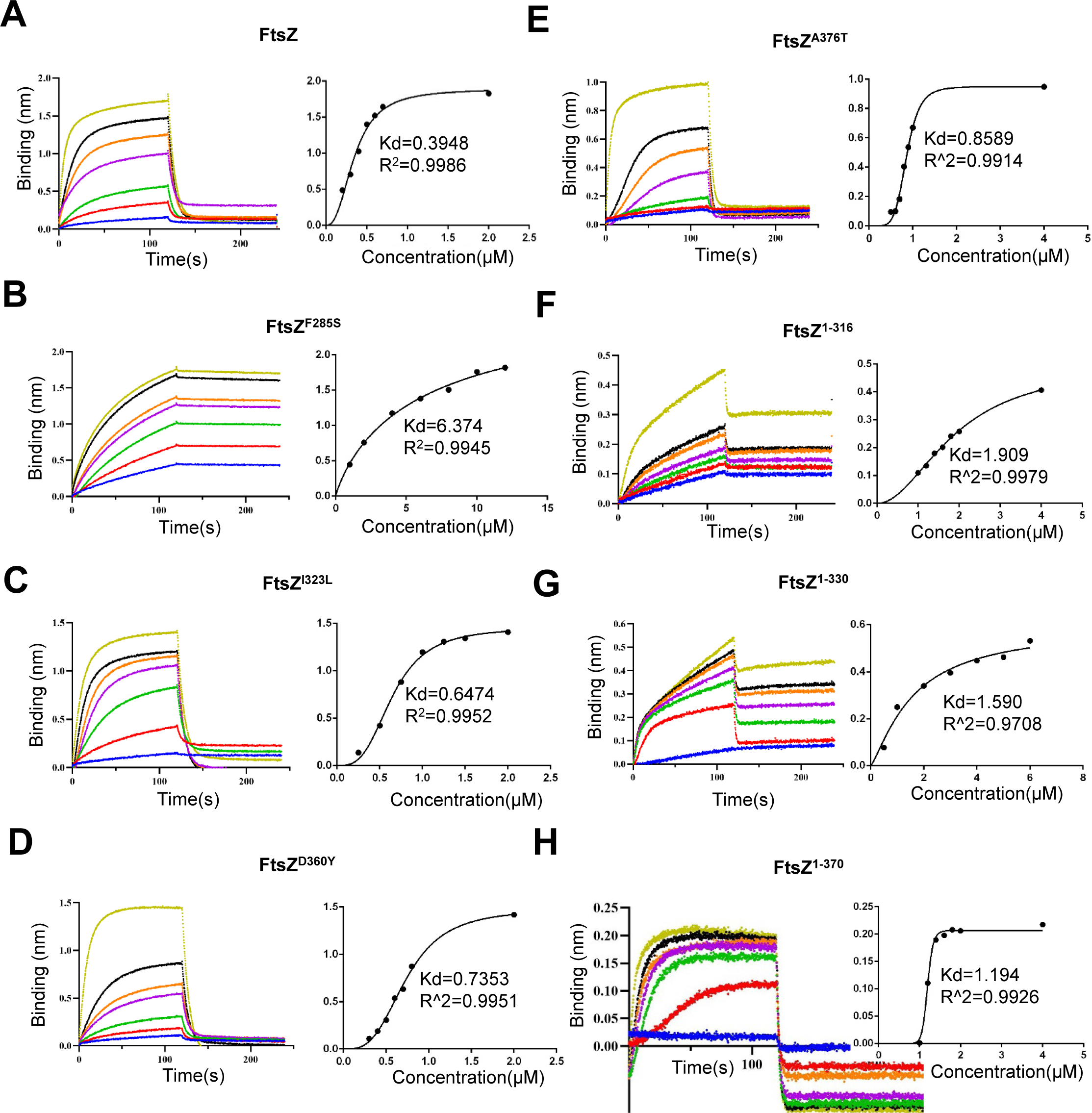
Multiple domains of FtsZ participate in the interaction with ZapC. Interaction between FtsZ and ZapC was assessed by biolayer interferometry assay. Details about the experimental set up and procedures are described in Materials and Methods. The concentration of 6×His-ZapC remained unchanged at 0.5 μM, but concentrations of FtsZ or its variants vary. Wild type FtsZ: 0.2 to 2 μM (A), FtsZ^F285S^: 1-12 μM (B), FtsZ^I323L^: 0.25-2 μM (C), FtsZ^D360Y^: 0.3-2 μM (D), FtsZ^A376T^: 0.5-4 μM (E), FtsZ^1-316^: 1-4 μM (F), FtsZ^1-330^: 0.5-6 μM (G), FtsZ^1-370^: 1-4 μM (H). Dissociation constant (Kd) of FtsZ or its variants for ZapC was determined by plotting the binding (nm) at the end of the association step with the concentration of proteins by GraphPad Prism.

### The globular domain and the CCTP of FtsZ bind to ZapC directly which are modulated by the CTL

The analysis of the ZapC resistant FtsZ mutants indicated that the globular domain, the CTL and the CCTP of FtsZ are all involved in the interaction with ZapC, with the globular domain playing a predominant role. To confirm this, we purified C-terminal truncated forms of FtsZ, including FtsZ^1-370^, FtsZ^1-330^ and FtsZ^1-316^ and measured their binding affinity to ZapC using the biolayer interferometry assay. These mutants lack the CCTP, most of the CTL and the CCTP, or the entire CTL and CCTP, respectively. As shown in Fig. 4F-H, the deletion of the CCTP of FtsZ (FtsZ^1-370^) reduced the binding affinity between FtsZ and ZapC about three fold, while simultaneously removing the CCTP and the CTL (FtsZ^1-316^ and FtsZ^1-330^), partially or entirely, further increased the Kd value (4-5 times compared to wild-type FtsZ and ZapC) (Table S4). Intriguingly, the binding kinetics of FtsZ^1-370^ to ZapC was similar to that of wild-type FtsZ, while FtsZ^1-316^ and FtsZ^1-330^ displayed a different binding profile (Fig. 4F-G). These results confirm that the globular domain of FtsZ binds to ZapC and both the CCTP and CTL affect the interaction.

To test if the CCTP and the CTL of FtsZ bind to ZapC directly, we purified SUMO-FtsZ and SUMO fusions containing only the linker region (FtsZ^316-370^), the CCTP (FtsZ^370-383^), or both (FtsZ^316-383^), and measured their binding to ZapC by biolayer interferometry assays. As shown in Fig. 5A-E, ZapC bound to both full length FtsZ and the SUMO fusions containing the CCTP but not the SUMO tag alone or the fusion carrying the CTL. However, the Kd value of the fusion containing the CTL and CCTP (SUMO-FtsZ^316-383^) was 6-7 times lower than that of full length FtsZ for ZapC (10 μM vs 1.5 μM), and was similar to the binding affinity of the fusion containing just the CCTP (SUMO-FtsZ^370-383^) for ZapC (Table S5). Interestingly, in the BTH assay we detected an interaction signal between ZapC and the fusion containing the CTL and the CCTP (FtsZ^316-383^), but the fusion containing only the CTL (FtsZ^316-370^) or only the CCTP (FtsZ^370-383^) did not display an interaction with ZapC (Fig. 5F), suggesting that although there is an interaction between ZapC and the CCTP of FtsZ *in vitro*, the presence of the CTL significantly enhances the interaction *in vivo*. Taken together, these results indicate that the CCTP of FtsZ, but not the CTL, binds to ZapC directly, however, the CTL modulates the binding.

**Figure 5.**
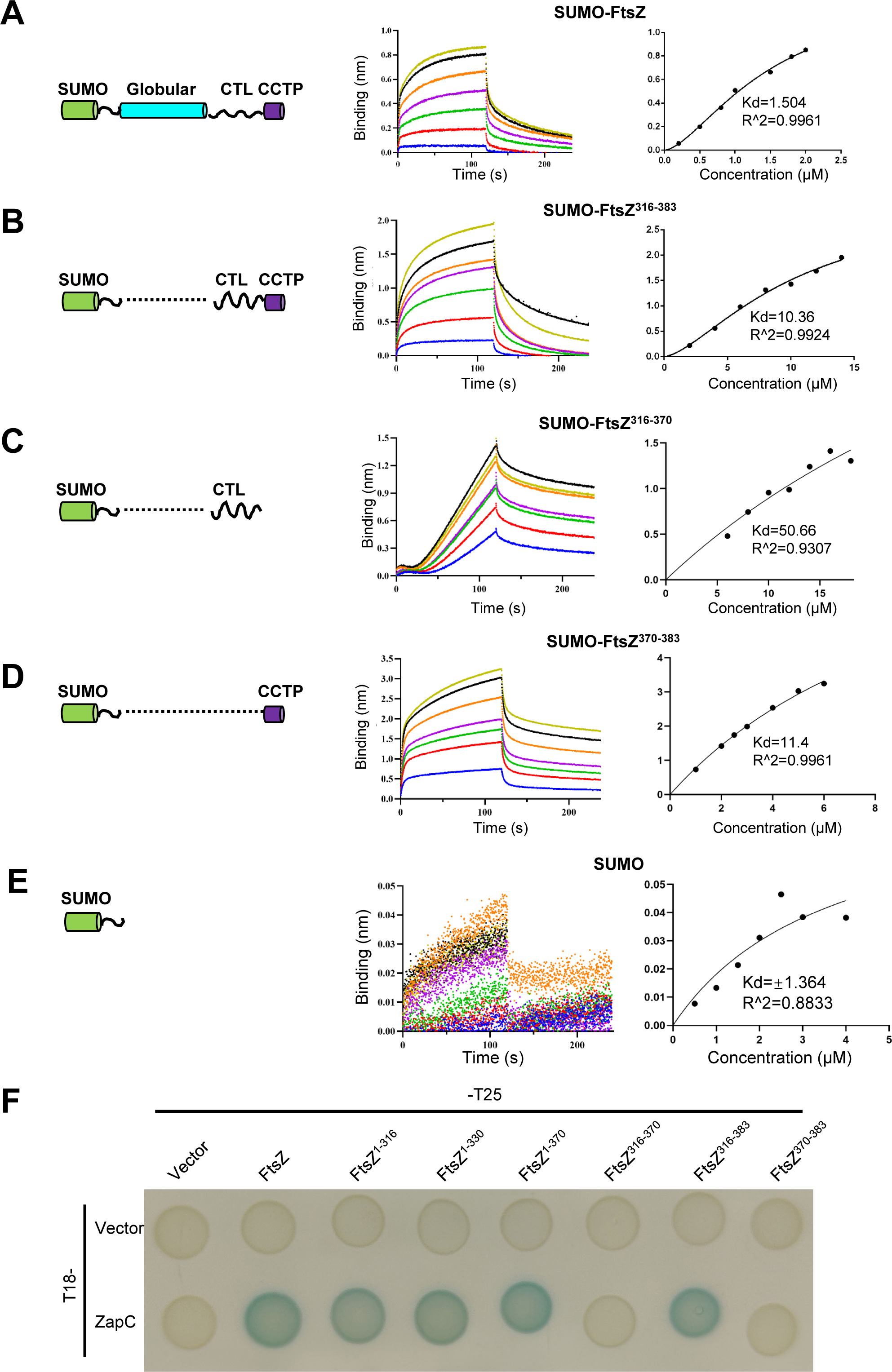
The CCTP of FtsZ binds to ZapC directly and is facilitated by the CTL of FtsZ. (A-E) Interaction between FtsZ, or its CCTP, and ZapC was determined by biolayer interferometry assay. The SUMO fusions with different parts of FtsZ is illustrated on the left and the results are shown on the right in each panel. Details about the experimental set up and procedures are described in Materials and Methods. The SUMO fusions remain unchanged at 0.5 μM and the concentration of ZapC varied according to different fusions. SUMO-FtsZ (A), SUMO-FtsZ^316-383^ (B), SUMO-FtsZ^316-370^ (C), SUMO-FtsZ^370-383^ (D) or SUMO (E). (F) The binding of the CCTP of FtsZ to ZapC requires the linkage to the CTL of FtsZ in bacterial two hybrid assay. The assay was performed as in Fig. 2B.

### Structural model of the FtsZ-ZapC complex indicates that ZapC binds to the globular domain and the CCTP of FtsZ

To further analyze the interaction between FtsZ and ZapC, we employed AlphaFold 3 to predict the structure of the FtsZ-ZapC complex (54). Strikingly, the structural model for the complex indicates that ZapC binds to both the globular domain and the CCTP of FtsZ but not the CTL (Fig. 6A), highly compatible with our genetic and biochemical results. In the first interaction site, the very conserved motif ^375^PAFLRK^380^ in the CCTP of FtsZ binds to the N-terminal pocket of ZapC largely via hydrophobic interactions. This motif forms a β-strand and binds to one of the β-strands in the N-terminal domain of ZapC, leading to the extension of the β-sheet. Analysis of the binding mode of the CCTP to ZapC in the model reveals a number of key interactions, including between residues L24, F30 and E72 of ZapC, and residues F377, L378 and K380 in the CCTP of FtsZ (Fig. 6B). Consistent with our results, substitutions in the CCTP (A376T, F377Y, K380M) provide resistance to ZapC overexpression *in vivo* and A376T displayed reduced ZapC binding to FtsZ *in vitro*. Notably, substitutions L22P and E72G that are close to the pocket of ZapC have been reported to disrupt its interaction with FtsZ (30). The second interaction site is largely consists of the loop region (^89^KPQMPKSW^96^) connecting the N-terminal and C-terminal domains of ZapC and a hydrophobic pocket on the surface of FtsZ, which consists of the region from I244-L254 and residue F285 (Fig. 6C). Residues P90, M92, K94 and W96 in the loop region of ZapC interact with the pocket of FtsZ. Notably, the side chain of M92 of ZapC appears to insert into the hydrophobic pocket by interacting with I244, L249 and L254 of FtsZ, and W96 interacts with F285 of FtsZ. The K94D mutation in ZapC was reported to prevent ZapC from binding to FtsZ in a previous study (47) and we found that the F285S/Y mutation in FtsZ greatly reduced the interaction between FtsZ and ZapC, lending support to the accuracy of the model. Thus, the structural model of the FtsZ-ZapC complex matches nicely with previous and current genetic and biochemical analysis of the interaction between FtsZ and ZapC.

**Figure 6.**
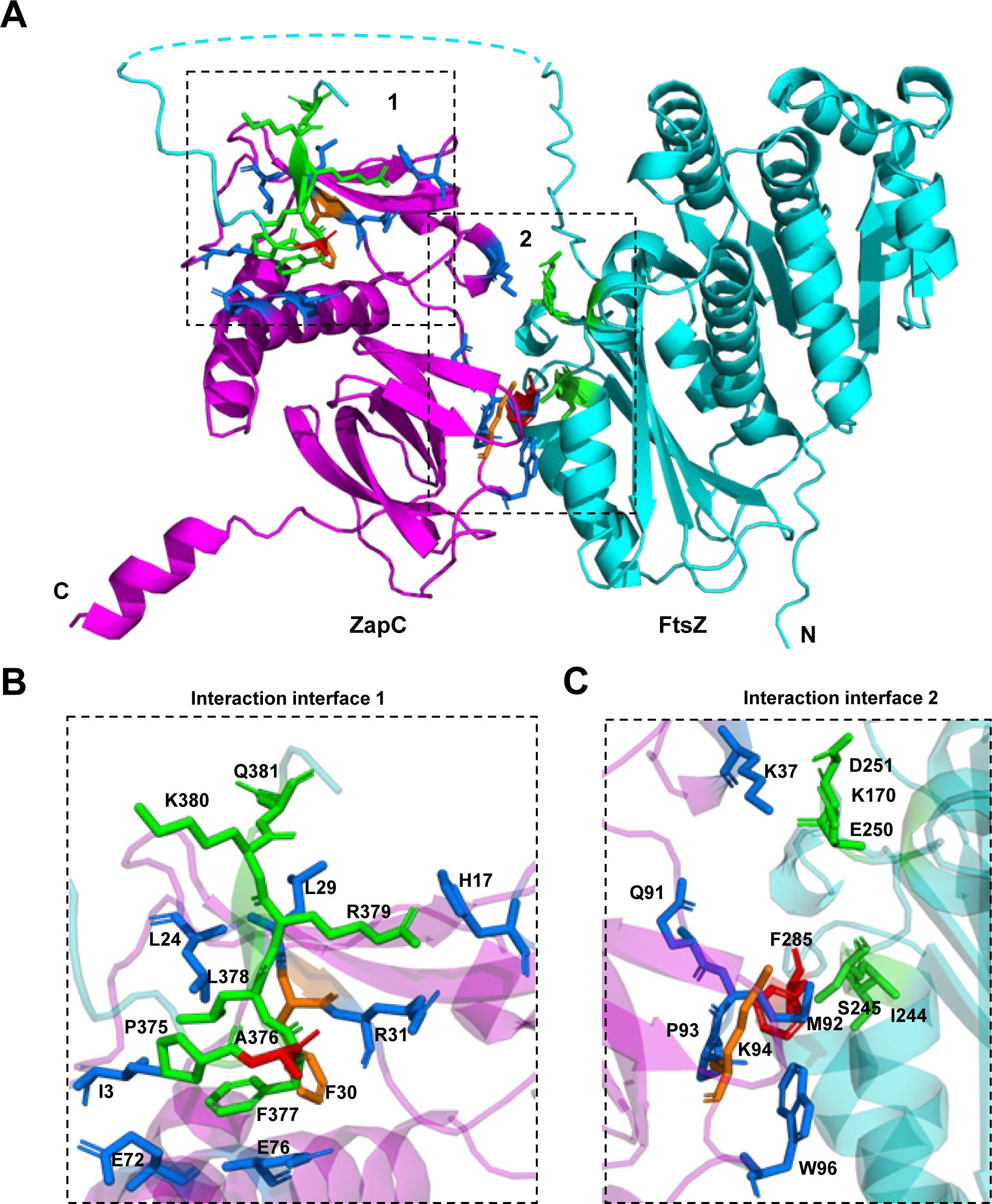
AlphaFold 3 structural model of the FtsZ-ZapC complex. The AF3 structure model reveals two interaction sites between FtsZ (cyan) and ZapC (magenta). The first interaction interface involves the CCTP of FtsZ and a pocket in the N-terminal domain of ZapC. The second interaction interface involves a hydrophobic pocket of FtsZ and the loop region connecting the N- and C-terminal domains of ZapC. Residues involved in the interactions are shown in stick and colored green (FtsZ) and blue (ZapC), respectively. Residues characterized in detail in previous studies and in this study were highlighted red or orange.

### Mutations in the two putative interaction interfaces of ZapC disrupt its interaction with FtsZ

The second putative interaction interface between FtsZ and ZapC (between the loop region of ZapC and the hydrophobic pocket of FtsZ near F285) is well supported by genetic and biochemical data, so we did not further investigate it. To validate the first interaction site from the structural model of the FtsZ-ZapC complex, F30 in the hydrophobic pocket of ZapC was mutated to aspartate and the mutant tested *in vivo* and *in vitro*. ZapC^F30D^ could not block cell division when it was overexpressed from a plasmid (Fig. 7A), similar to the well-characterized ZapC^K94D^ mutant in the second putative interaction site, suggesting that the mutation weakens the interaction with FtsZ. Consistent with this loss of toxicity, ZapC^F30D^ (fused with GFP) was evenly distributed in the cytoplasm and no longer co-localized with ZapA-mCherry (Fig. 7B). Moreover, ZapC^F30D^ did not interact with FtsZ in the BTH assay (Fig. 7C) and Western blot analysis showed that it was expressed at comparable levels as wild type ZapC (Fig. S5). Thus, mutation in the first interaction interface greatly reduced the interaction between ZapC and FtsZ *in vivo*.

**Figure 7.**
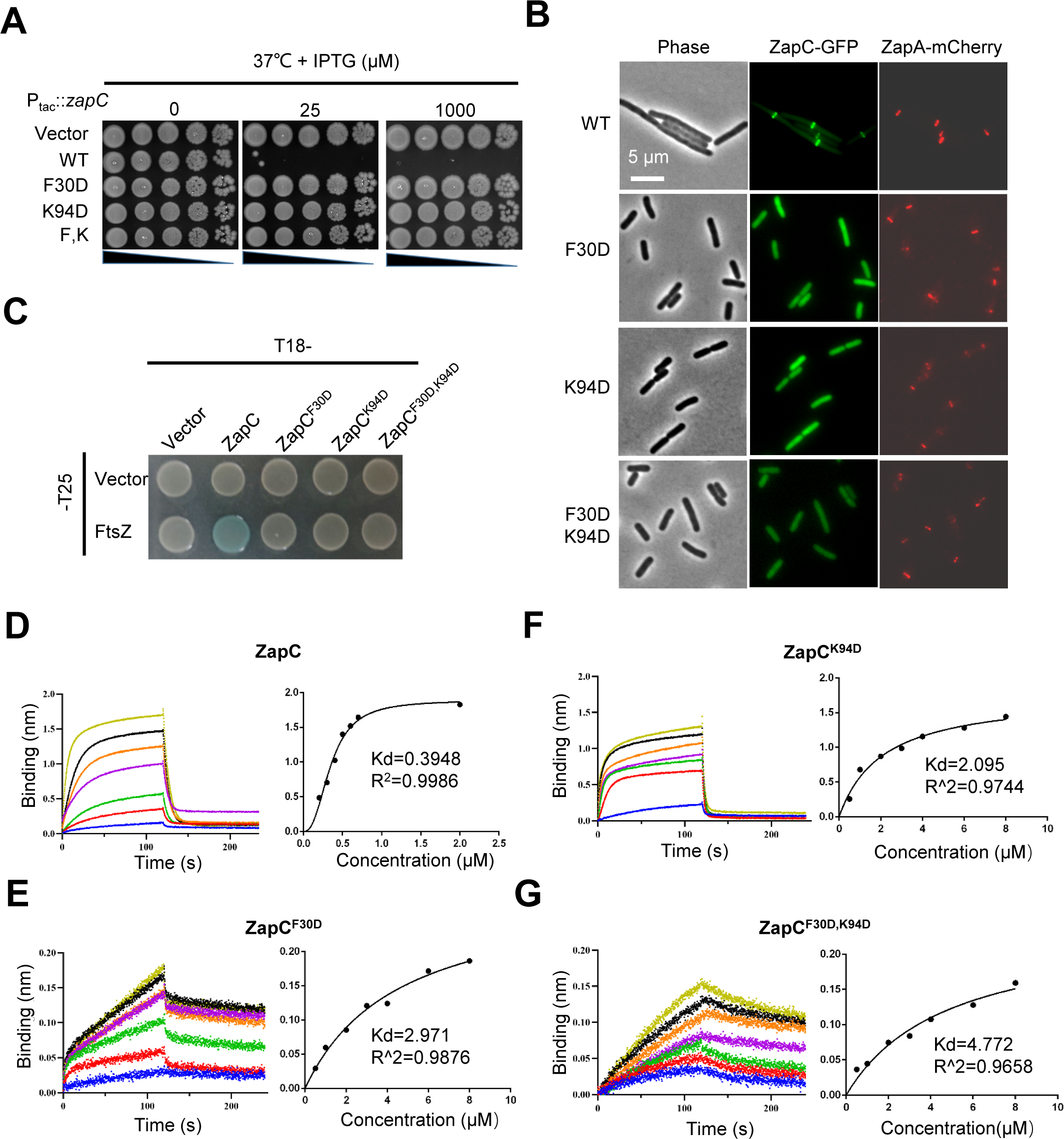
Mutations in the two putative interaction interfaces of ZapC synergize to disrupt its interaction with FtsZ. (A) Mutations in the two putative interaction interfaces of ZapC reduce its toxicity. Plasmid pSD320 (pEXT22, P_tac_::*zapC*) or its derivatives was transformed into strain LYA4 (TB28, *zapA-mCherry*) at 37°C with kanamycin. The next day, a single colony of each resulting strain was subject to spot tests. Plates were incubated at 37°C for 24 hours and photographed. (B) Mutations in the two putative interaction interfaces of ZapC reduce the midcell localization of ZapC. Overnight cultures of LYA4 (TB28, *zapA-mCherry*) carrying plasmid pLY44 (pDSW210, P_206_::*zapC-l60-gfp*) or its derivatives were diluted 1:100 in fresh LB medium with antibiotics and cultured at 37°C. 3 hours later the cultures were diluted 1:10, IPTG was added to a final centration of 100 μM to induce the expression of the ZapC-GFP. 2 hours post induction cells were immobilized on 2% agarose pads and photographed. Scale bar, 5 μm. (C) Mutation in either the first or the second interaction site of ZapC blocks its interaction with FtsZ in the bacterial two hybrid test. The test was performed as Fig. 2B. (D) Mutation in either the first or the second interaction site of ZapC reduces its binding affinity for FtsZ in biolayer interferometry assay. Details about the experimental setup and procedures were described in Materials and Methods. The concentration of 6×His-ZapC or its mutants was 0.5 μM and increasing concentration of FtsZ was added to the reaction system and binding kinetics monitored. Dissociation constant (Kd) of FtsZ for ZapC or its variants was determined by plotting the binding (nm) at the end of the association step with the concentration of proteins by GraphPad Prism.

To further confirm the involvement of the first hydrophobic pocket of ZapC in its interaction with FtsZ, we purified ZapC^F30D^ and tested its interaction with FtsZ by the biolayer interferometry assay. His-tagged ZapC or its variants were attached to the sensors and untagged FtsZ was employed to assess the binding. As shown in Fig. 7D-E, ZapC^F30D^ and ZapC^K94D^ still bound to FtsZ, but their binding affinities for FtsZ were 5-7 times lower than that of wild-type ZapC (Table S6). Moreover, the double mutant further reduced the binding to FtsZ, with a Kd value 12 times higher than that of wild type ZapC with FtsZ (Fig. 7G) (Table S6). Taken together, these results suggest that both interaction interfaces are important for ZapC binding to FtsZ and mutations in any one greatly reduces ZapC’s interaction with FtsZ.

## Discussion

Formation of a condensed and stable Z ring is the first step of bacterial cytokinesis. *E. coli* and many closely related bacteria employ Zap proteins, which are capable of crosslinking FtsZ filaments, to facilitate Z ring formation and stabilization (27, 34). Interestingly, oligomeric ZapA or ZapD crosslinks FtsZ filaments by binding to the N-terminal tail or CCTP of adjacent filaments, but for ZapC, a monomer, it was not yet clear how it crosslinks filaments. In this study, we find that ZapC binds to the globular domain and the CCTP of FtsZ via two interaction sites. These findings suggest a model in which a ZapC monomer crosslinks adjacent FtsZ filaments by binding to the globular domain of an FtsZ molecule in one filament and the CCTP of an FtsZ molecule in another filament (Fig. 8). Moreover, the CTL of FtsZ can modulate the binding of the CCTP to ZapC. Given that many FtsZ regulatory proteins bind to the globular domain and the CCTP of FtsZ simultaneously, this finding suggests that the CTL, albeit being intrinsically disordered and not conserved in amino acid sequence, plays a role in the regulation of the interaction between FtsZ and its binding partners.

**Figure 8.**
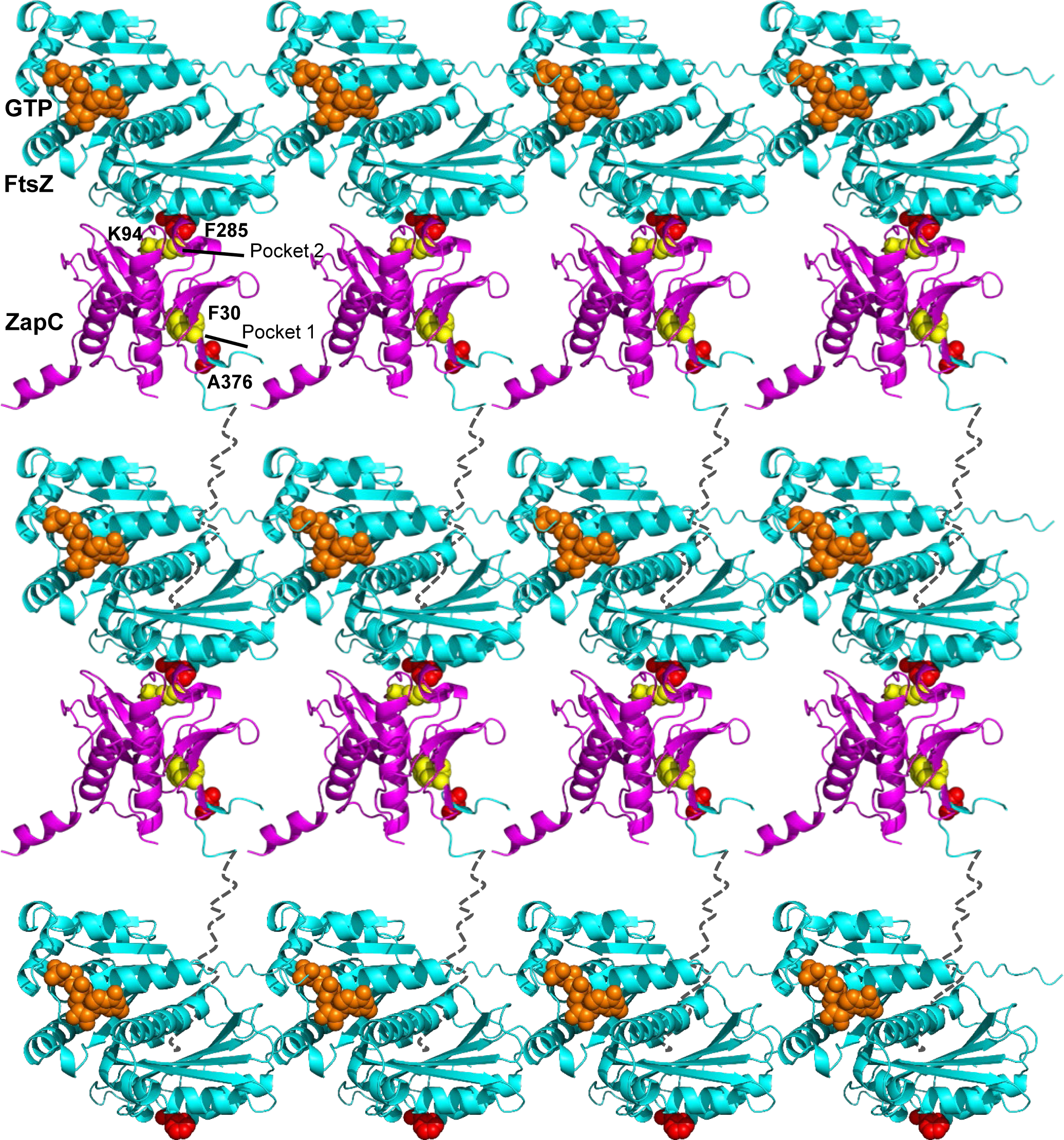
A working model for the crosslinking activity of ZapC on FtsZ filaments. ZapC (magenta) grabs the CCTP of FtsZ within a filament through its N-terminal hydrophobic pocket. Meanwhile, the loop connecting the N- and C-terminal domains of ZapC binds to the globular domain of FtsZ (cyan) (F285 interface, red) of an adjacent filament. These two bindings, allows monomeric ZapC to crosslink FtsZ filaments efficiently. Note that the CTL (black dash line) of FtsZ can modulate the binding of the CCTP to ZapC.

Although all the Zap proteins (except for ZapB) can crosslink FtsZ filaments, their working mechanisms are likely distinct. ZapA forms tetramers so that it is conceivable that it crosslink FtsZ filaments if each subunit binds to one FtsZ filament (35, 39). ZapD forms dimers with each subunit containing a binding site for the CCTP of FtsZ (45). Therefore, it is also not surprising that it can crosslink adjacent FtsZ filaments. However, unlike ZapA and ZapD, ZapC is a monomer (47) and how it crosslinks FtsZ filaments has been mysterious. Previous studies found that ZapC contains two pockets, mutations in which disrupt ZapC binding to FtsZ (47). Thus, it was proposed that each of the two pockets bind to different parts of the globular domain of FtsZ to crosslink FtsZ filaments (47). However, another study observed that ZapC has a tendency to form dimers at high concentration and detected a weak binding of ZapC to the CCTP of FtsZ (48), raising a possibility of ZapC dimers crosslink FtsZ filaments by binding to the CCTP.

To determine the exact mechanism by which ZapC crosslinks FtsZ, we isolated FtsZ mutations that confer resistance to ZapC overexpression. Several lines of evidence suggest that the corresponding residues are involved in the interaction between FtsZ and ZapC. First, the midcell localization of ZapC-GFP was reduced to varying degrees by these mutations. Second, these FtsZ mutations reduced the interaction between FtsZ and ZapC in the BTH assay. Third, these mutations markedly weaken the interaction between FtsZ and ZapC *in vitro* as determined by a sedimentation assay, negative staining electron microscopy and biolayer interferometry experiments. Intriguingly, instead of clustering together, these mutations are scattered and located in the globular domain, the CTL and the CCTP of FtsZ, suggesting that all these regions of FtsZ are involved in its interaction with ZapC. This explains why the single mutations could not completely eliminate the toxicity of ZapC or localization of ZapC. For example, the F285S mutation in the globular domain and the A376T mutation in the CCTP could each significantly reduce the toxicity and localization of ZapC, but a combination of them resulted in greater resistance and elimination of ZapC localization. Although mutations in the CTL of FtsZ affected the interaction between FtsZ and ZapC in the genetic and biochemical tests, subsequent analysis of the interaction between FtsZ and ZapC using truncated FtsZ mutants found that the globular domain and the CCTP of FtsZ directly bind to ZapC, whereas the CTL affects the binding indirectly. An AlphaFold 3 structural model of the FtsZ-ZapC complex indicates that ZapC binds to the globular domain and the CCTP but not the intrinsically disordered CTL, lending support for our findings. Moreover, introduction of mutations into the putative binding sites for FtsZ on ZapC eliminated its toxicity and prevented it from localizing to the division site. Taken together, these results provide compelling evidence that ZapC binds to both the globular domain and the CCTP of FtsZ, reconciling the contradictory observations reported by other research teams previously (ZapC binding to the globular domain but not CCTP of FtsZ *vs* ZapC binding to the CCTP of FtsZ).

Examination of the binding mode between ZapC and FtsZ reveals that the N-terminal hydrophobic pocket of ZapC binds to the CCTP of FtsZ, whereas the loop region connecting its two subdomains binds to a pocket near residue F285 in the globular domain of FtsZ. The F285S mutation results in a much stronger reduction in the interaction than mutations in the CCTP, suggesting that the interaction between the loop region of ZapC and the globular domain of FtsZ is stronger than the interaction involving the hydrophobic pocket of ZapC and the CCTP of FtsZ. Moreover, the CCTP alone binds to ZapC with a rather weak affinity, which is consistent with a previous report (48). It has been noted that CCTP binding proteins bind to full length FtsZ with high affinity, usually in the nanomolar range, but display low affinity for the CCTP alone. We previously showed that polymerization of FtsZ converted FtsZ to a multivalent ligand, such that ZipA and SlmA bind to FtsZ filaments/oligomers with high affinity due to avidity but FtsZ monomers bind with low affinity (55). This situation may also apply to ZapC. We suspect that ZapC initially grabs the CCTP of FtsZ in a filament and then binds to the globular domain of FtsZ on an adjacent filament to crosslink them. It is not clear how ZapC avoids binding to both sites on the same FtsZ molecule or binds two FtsZ molecules in the same filament concurrently. Of course, this is true of all Zap proteins.

The CCTP of FtsZ is widely conserved and absolutely critical for FtsZ function in cell division (2, 5). In *E. coli*, a number of proteins have been found to bind to it, including the membrane anchors for FtsZ filaments (FtsA and ZipA)(10, 56–58), antagonists of FtsZ polymerization (MinC and SlmA)(12–14), cross-linker of FtsZ filaments (ZapD) and proteases that degrade FtsZ (ClpXP)(45, 59). It is also the hub for FtsZ binding proteins in other bacteria. Thus, it is perhaps not surprising that ZapC also binds to it. The addition of ZapC to the growing list of CCTP binding proteins further underscores the importance and flexibility of this short motif in governing the function of FtsZ. Previous studies have found that the CCTP of FtsZ is a classical conserved short linear motif (SLiM) embedded in an intrinsically disordered region (IDR). Consistent with well characterized SLiMs it acquires various secondary structures upon binding to partner proteins (2). For example, when bound to ZipA or FtsA it is partially folded into a helix (57, 58), while in the complex with SlmA tetramer bound with DNA, it adopts an unusual extended loop-like conformation (14). The structural model of the FtsZ-ZapC complex suggests that the CCTP is partially folded into a β strand, leading to the extension of the β sheets of ZapC. Although further structural analysis of the ZapC-CCTP complex is necessary to confirm this conformation, this high adaptability of the CCTP makes it a perfect example to study SLiMs and IDRs in prokaryotes.

Perhaps one of the most intriguing observations in this study is that single substitutions in the CTL of FtsZ could significantly affect its interaction with ZapC. Initially, we thought the CTL was directly involved in the binding of ZapC, however, subsequent analysis found that it likely exerts its effect on the interaction through its influence on the CCTP. Strikingly, we found that the CTL’s impact on the interaction between the CCTP and ZapC is not an exception, it also strongly affects the binding of the CCTP to many other FtsZ binding proteins. As shown in Fig. 9, neither the CTL nor the CCTP displayed a binding signal with FtsA, SlmA or MinC, all of which have been confirmed to bind to the CCTP, in the BTH assay, but the fusion containing both the CTL and CCTP interacted with these proteins strongly. This suggests that the CTL strongly affects the property or configuration of the CCTP and modulates its binding to FtsZ binding partners. Previous studies have showed that although the CTL is hypervariable across FtsZ proteins from diverse bacterial species, including length, amino acid composition, and sequence, its presence is absolutely essential for proper FtsZ function (15–18). From *E. coli* to *Caulobacter crescentus* and *Bacillus subtilis*, the absence of the CLT resulted in nonfunctional FtsZ variants, leading to the assembly of aberrant FtsZ structures *in vivo* and *in vitro*. Replacement of the CTL with IDRs with similar property usually generates partially functional FtsZ proteins (15, 16), suggesting that amino acid composition is probably not important for its function. However, in contrary to this early suggestion, a recent study found that the sequence patterns of the CTLs of FtsZ are non-random (18). Instead, they are evolutionary conserved and dictate the conformational properties of CTLs, mediating autoregulatory interactions between covarying regions within FtsZ (18). As a result, disruption of these patterns led to abnormal assembly of FtsZ *in vivo* and *in vitro*, and disruption of cell division *in vivo* (*18*). Here, we provided direct evidence that the CTL modulates FtsZ interaction with ZapC and other CCTP-binding proteins. It is still difficult to imagine how the CTL affects FtsZ binding to its partner proteins because it was not resolved in any of the available structures of FtsZ, regardless of monomeric, oligomeric or in complexes with its binding partners. Perhaps the mutations alter the conformational properties of the CTL, such as the radius of gyration and the end-to-end distance (measures of global ensemble dimensions), asphericity (a measure of ensemble shape), transient secondary structure (a measure of local structural acquisition) and inter-residue distances (a measure of specific ensemble dimensions) (60), such that the affinity between FtsZ and ZapC is changed. It will be interesting to elucidate the mechanism by which the CTL modulates its binding to ZapC as well as other partner proteins in future investigations.

**Figure 9.**
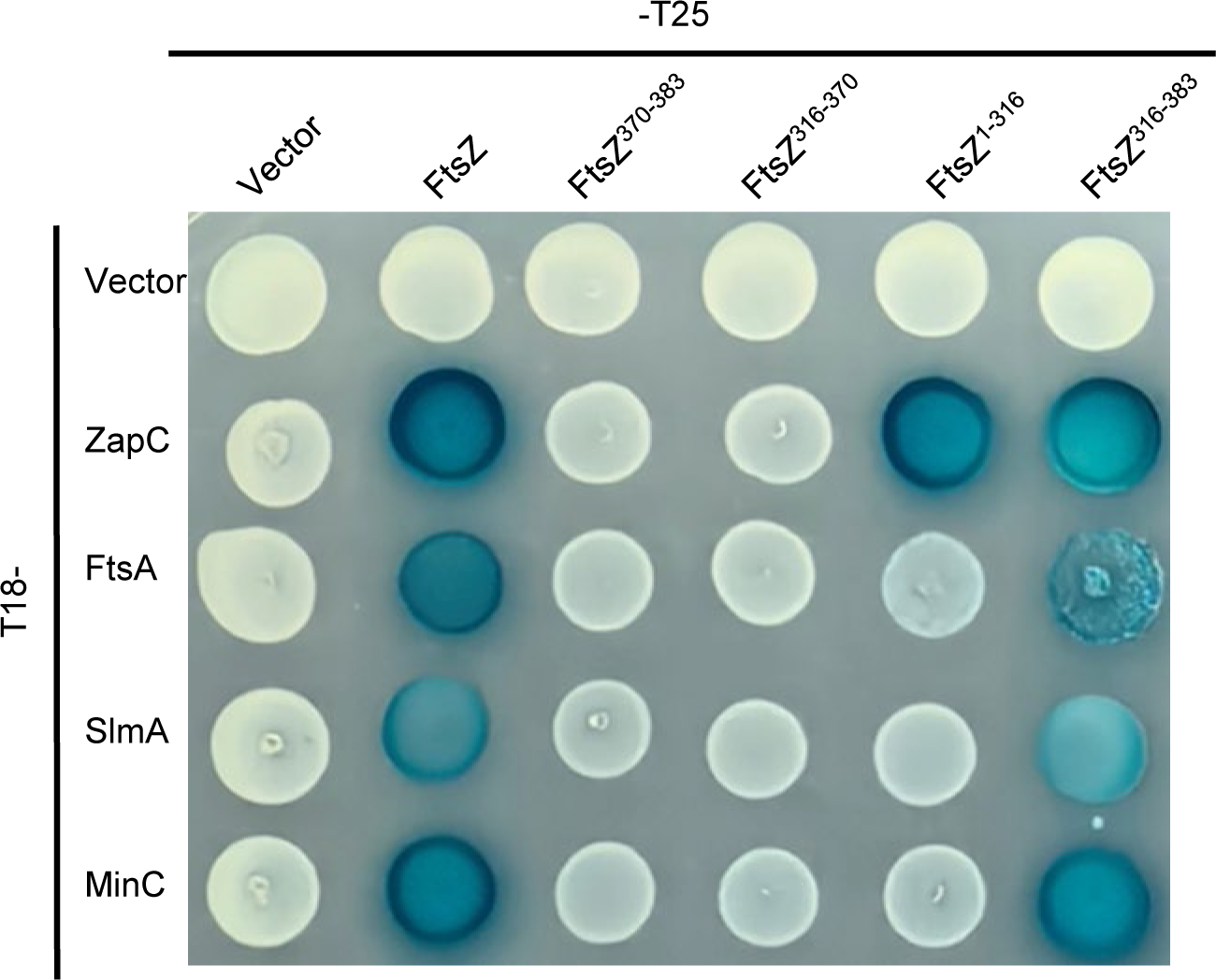
The CTL of FtsZ modulates its binding to its partners as indicated by bacterial two hybrid assay. Plasmids pairs expressing the indicated fusions were transformed into strain LYA1 (BTH101, *ftsA*^R286W^). The next day a single transformant of each resulting strain was resupended in 1 mL LB medium and 2 μL of each culture was spotted on LB plates containing antibiotics, 100 μg/mL X-gal and IPTG. Plates were incubated at 30°C for about 40 hours before photographing.

## Materials and methods

### Media, bacterial strains, plasmids and growth conditions

Cells were grown in LB medium (1% tryptone, 0.5% yeast extract, 0.5% NaCl and 0.05 g/L thymine) at indicated temperatures. When needed, antibiotics were used at the following concentrations: ampicillin = 100 μg/mL; kanamycin = 25 μg/mL; and chloramphenicol = 20 μg/mL. Strains, plasmids and primers used in this study are listed in Table S7-9, respectively. Construction of strains and plasmids is described in detail in Supplemental Information with the primers listed in Table S9.

### Construction of the FtsZ mutant library and screen for FtsZ mutants resistant to ZapC overexpression

*Construction of the FtsZ mutant library*. *ftsZ* was subjected to random PCR mutagenesis using the primer FtsZ-F and FtsZ-R and the plasmid pBANG112 (p15A, *ftsZ*) as a template. Purified PCR product was digested with SacI and EagI and ligated into pBANG112 digested with the same enzymes. Ligation product was transformed into JS238 competent cells and plated on LB plates with ampicillin and glucose. About 40,000 transformants were pooled together and plasmids were isolated and stocked.

#### Screen for FtsZ mutants resistant to ZapC overexpression

The mutagenized pBANG112 (pBANG112^M^) library was transformed into strain S17/pKD3C [W3110, *ftsZ*^0^/pSC101^ts^, *ftsZ*] harboring plasmid pSD320 (pEXT22, P_tac_::*zapC*). Transformants were selected on LB plates with antibiotics and 60 µM IPTG at 42°C. Strain S17/pKD3C could not grow at 42 °C because plasmid pKD3C was not able to replicate. Transformants that grew on the selective plates thus contained pBANG112 derivatives expressing FtsZ mutants that could compensate for the depletion of FtsZ and confer resistance to the toxicity of ZapC overexpression. 15 transformants were randomly picked and restreaked on the selective plates for purification. Plasmids were isolated from these 15 transformants and retransformed into strain S17/pKD3C carrying pSD320 to confirm the resistance. *ftsZ* in these 15 suppressing plasmids was sequenced and 13 of them harbored one or more mutations in *ftsZ* (Table S1). Each mutation was then introduced into the parental plasmid pBANG112 using site-directed mutagenesis and tested for resistance to ZapC overexpression toxicity to identify the causative mutation. A total of 10 mutations in FtsZ were identified to confer ZapC-resistance (E147G, F285S/Y, E322K, I323L, N359Y, D360Y, A376T, F377Y and K380M) (Table S2).

### BTH assay

To detect the interaction between FtsZ or its mutants and ZapC, appropriate plasmid pairs were co-transformed into BTH101 or the strain LYA1 (BTH101, *ftsA\**). The next day, single colonies were resuspended in 1 ml of LB medium, and 2 μL of each aliquot was spotted on LB plates containing ampicillin, kanamycin, 40 μg/mL X-gal and 10 μM IPTG. Plates were incubated at 30°C overnight or longer before imaging.

### Co-localization of ZapC-GFP and ZapA-mCherry

#### Effect of FtsZ mutation on the co-localization of ZapC

*GFP*. Overnight cultures of LYA6 [TB28, *zapA*-*mCherry ftsZ*^0^] carrying plasmid pLY44 (pDSW210, P_206_::*zapC*-*l60*-*gfp*) and pBANG112 (p15A, *ftsZ*) or its derivatives were diluted 1:100 in fresh LB medium with antibiotics, grown at 30 °C for 3 h. Cells were then diluted 1:10 in fresh LB medium and IPTG was added to a final concentration of 50 µM. 2.5 hours post the addition of IPTG, cells were immobilized on 2% agarose pad for photograph.

#### Effect of ZapC mutation on the co-localization of ZapC-GFP and ZapA-mCherry

Overnight cultures of LYA6 [TB28, *zapA*-*mCherry ftsZ*^0^] carrying plasmid pBANG112 ( p15A, *ftsZ* ) and pLY44 (pDSW210 , P_206_::*zapC*-*l60*-*gfp*) or its derivatives were diluted 1:100 in fresh LB medium with antibiotics, grown at 30 °C for 3 h. Cells were then diluted 1:10 in fresh LB medium and IPTG was added to a final concentration of 50 µM. 2 hours post the addition of IPTG, cells were immobilized on 2% agarose pad for photograph.

### Western blot

To measure the level of ZapC-GFP mutants, overnight cultures of W3110 harboring the respective plasmids were diluted 1:100 in LB medium with kanamycin and 30 µM IPTG. After growth at 37°C for 2 hours, OD_600_ of each culture was measured and samples were taken for western blot. Cells were collected, resuspended in SDS-PAGE sample buffer and kept at 95°C or 10 min before they were loaded onto the SDS-PAGE gel for analysis. Anti-GFP antibody and Anti-FtsZ serum were used at a dilution of 1/10000.

### Protein purification

FtsZ and its mutants were expressed in *E. coli* strain BL21 (DE3) harboring plasmid pLY17 (pE-SUMO-Amp, P_T7_::*6×his-sumo-ftsZ*) or its derivatives. Briefly, bacteria were grown at 37 °C in 1 L LB medium supplemented with 100 µg/mL ampicillin to an OD_600_ of 0.6. Isopropyl β-D-1-thiogalactopyranoside (IPTG) was then added to the culture to induce the expression of proteins. 3 hours post the induction, cells were collected by centrifugation, resuspended in 20 ml lysis buffer (25 mM Tris-HCl [pH 7.5], 300 mM NaCl, 5% glycerol, 0.1 mM DTT and 20 mM imidazole) and lysed by sonication. The lysates were centrifuged at 10,000 rpm for 10 min at 4°C to remove cell debris. The supernatants were loaded onto pre-equilibrated Ni-NTA resin (Qiagen). The column was washed once with high salt wash buffer (25 mM Tris-HCl [pH 7.5], 500 mM NaCl, 5% glycerol, 0.1 mM DTT and 20 mM imidazole) and then once with the same buffer except with the imidazole concentration increased to 50 mM. The bound protein was eluted with elution buffer (25 mM Tris-HCl [pH 7.5], 500 mM NaCl, 5% glycerol, 0.1 mM DTT and 250 mM imidazole) and analyzed with SDS-PAGE. The peak fractions were pooled and dialyzed against the storage buffer (50 mM Tris-HCl [pH 7.5], 300 mM NaCl, 5% glycerol and 0.1mM DTT) overnight and stored at −80°C until use.

Expression and purification of ZapC or 6×His-ZapC and its mutants were similar to the procedure of FtsZ using plasmids pSD323 (pQE80, P_lac_::6×His-ZapC) and pLY28 (pE-SUMO-Amp, P_T7_::*6×his-sumo-zapC*) or its derivatives. Because ZapC contains multiple cysteine residues and tend to form aggregates, the final concentration of DTT in the dialysis buffer is 1 mM. To remove the 6×His-SUMO tag, the purified proteins were mixed with Ulp1 protease, incubated at 4°C overnight or at 30°C for 1 hour. The released tag and protease were removed by passing the mixtures through the pre-equilibrated Ni-NTA resin. Untagged FtsZ or ZapC was collected in the flow through and wash fractions, concentrated and stored at −80°C until use.

### Co-sedimentation assay

FtsZ polymerization reactions were performed in 50 μL (final volume) of Pol buffer (50 mM HEPES-KOH, 50 mM KCl, 10 mM MgCl_2_, pH 6.8) at room temperature. After the addition of FtsZ (5 μM), GTP or GDP was added to a final concentration of 1 mM. 5 min later, ZapC or CaCl_2_ was added to 5 μM or 1 mM, respectively, and then the mixtures were subjected to high-speed centrifugation (100, 000 rpm) for 12 min at 25°C in a BACKMAN OPTIMA MAX XP ultracentrifuge with an MLA 130 rotor. Pellets were resuspended in 50 μL of buffer, and equal amounts of pellet and supernatant fractions were loaded onto SDS-PAGE gels for analysis.

### Transmission electron microscopy

To visualize the effect of ZapC on the assembly of FtsZ and its mutants, the polymerization reaction (50 μL, final volume) was carried out in Pol buffer (50 mM HEPES-OH (pH 6.8), 50 mM KCl, 10 mM MgCl_2_) in the presence or absence of ZapC at room temperature. FtsZ and ZapC was added to a final concentration of 5 μM and 1 mM GTP or GDP was added to initiate FtsZ polymerization. After incubation for 5 min, 15 μL aliquot was applied to a carboncoated copper grid (100-mesh). 1 min later, the excess solution was absorbed with filter paper, and stained with 15 μL 1% uranyl acetate for 1 min. Grids were air-dried for more than 12h and then examined with a JEM 1400 plus transmission electron microscope (TEM) at 100 kV.

### Biolayer Interferometry assays

Biolayer Interferometry assays were performed with the Octet-Red96 BLI system at 30°C. To detect the binding of ZapC to FtsZ or its variants, 6×His-ZapC (0.5 µM) was loaded on the pre-equilibrated Ni-NTA biosensors for 3 minutes in 200 µL of 1×Pol buffer (50 mM HEPES-NaOH [pH 6.8], 50 mM KCl, and 10 mM MgCl_2_) and then washed with the same buffer for 30 seconds to remove any loosely bound protein. Binding of FtsZ or its mutants to the immobilized His-ZapC was monitored for 2 minutes with agitation at 1000 rpm followed by dissociation in the same buffer without proteins for 2 minutes. To detect the interaction between ZapC and different domains of FtsZ, SUMO fusions containing full length FtsZ, its linker or CCTP were immobilized on the Ni-NTA biosensors as above, binding of untagged ZapC to these SUMO fusions was then monitored for 2 minutes followed by dissociation. Data were automatically collected by the Octet-Red96 BLI system and analyzed with GraphPad Prism 6. To obtain the apparent dissociation constant value (Kd), the binding signals at the end of the association step were plotted against the protein concentrations using a one-site specific nonlinear regression fitting.

## Acknowledgments

We thank members of the Du lab, Chen lab and Lutkenhaus lab for comments and advice in preparing the manuscript. This study was supported by National Natural Science Foundation of China (grant 32070032 and 32270049 http://www.nsfc.gov.cn/), the Fundamental Research Funds for the Central Universities (grant 2042021kf0198), and the Young Top-notch Talent Cultivation Program of China to S.D., and Natural Science Foundation of Hubei Province (2025AFB238, http://kjt.hubei.gov.cn/) to Y.L..

## Supplemental Information

Figure S1-S5

Table S1-S9

